# A Fluorescence-Based Sensor for Calibrated Measurement of Protein Kinase Stability in Live Cells

**DOI:** 10.1101/2023.12.07.570636

**Authors:** Joseph W. Paul, Serena Muratcioğlu, John Kuriyan

**Author notes:** Correspondence to John Kuriyan.

## Abstract

Oncogenic mutations can destabilize signaling proteins, resulting in increased or unregulated activity. Thus, there is considerable interest in mapping the relationship between mutations and the stability of proteins, to better understand the consequences of oncogenic mutations and potentially inform the development of new therapeutics. Here, we develop a tool to study protein-kinase stability in live mammalian cells and the effects of the HSP90 chaperone system on the stability of these kinases. We monitor the fluorescence of kinases fused to a fluorescent protein relative to that of a co-expressed reference fluorescent protein. We used this tool to study the dependence of Src- and Raf-family kinases on the HSP90 system. We demonstrate that this sensor reports on destabilization induced by oncogenic mutations in these kinases. We also show that Src-homology 2 (SH2) and Src-homology 3 (SH3) domains, which are required for autoinhibition of Src-family kinases, stabilize these kinase domains in the cell. Our expression-calibrated sensor enables the facile characterization of the effects of mutations and small-molecule drugs on protein-kinase stability.

## Introduction

An initiating step in cancer is the unregulated activation of signaling proteins, such as protein kinases, by mutations that override negative regulation. The earliest identified viral-encoded oncogenes v-Src^1,2^, v-Raf^3^, and v-Abl^4,5^ are mutant forms of cellular protein kinases. These mutant kinases exhibit unregulated kinase activity in contrast to their normal cellular counterparts, which underlies their ability to transform healthy cells into cancerous ones^6–8^. This hyperactivity often comes at the expense of structural stability. For example, v-Src protein has a lower threshold for irreversible unfolding compared to endogenous, normal cellular c-Src^9^. The reduced stability and hyperactivity of oncogene-encoded proteins impedes high-level heterologous expression, particularly in bacteria, for in vitro studies.

The common ancestor of eukaryotes evolved specialized protein chaperone systems dedicated to protein kinases. Experiments in yeast revealed that two conserved chaperones, heat shock protein 90 (HSP90) and a co-chaperone, cell division cycle protein 37 (CDC37), bind to and preserve the activity of oncogenic v-Src but do not bind to c-Src^10^. When the genes encoding HSP90 or CDC37 are deleted, or when these proteins are inhibited pharmacologically in yeast, v-Src is rapidly degraded by the ubiquitin proteasome system, indicating that chaperones protect v-Src from degradation^10,11^. Studies in *C. elegans* and *D. melanogaster* established a physiologic function for the HSP90/CDC37 system in maintaining the mitogen-activated protein kinase signaling pathway. Genetic ablation of HSP90 or CDC37 in certain cell lineages phenocopies the developmental defects caused by mutations in a Raf kinase homolog or its upstream regulators, such as homologs of the epidermal growth factor receptor, Son-Of-Sevenless, and Ras GTPase^12,13^.

The HSP90/CDC37 complex directly interacts with many tyrosine and serine/threonine kinases in cells^14^. The dependence of these kinases on the HSP90 system can vary among closely related proteins and is not linked to any specific sequence motif^14^. For example, in vertebrate Raf-family kinases, ARAF and CRAF undergo rapid proteolytic degradation when cells are treated with HSP90 inhibitors, while these inhibitors do not affect BRAF levels^15^. Remarkably, a single, activating point mutation in BRAF, V600E, is sufficient to convert BRAF into an obligate client of HSP90^15^. The V600E mutation is a common driver of melanoma and other cancers^16^. Val 600 is located in the activation loop of the kinase domain and the hydrophobic valine sidechain stabilizes the activation loop in an autoinhibitory conformation^17^. Substitution of valine by the charged residue glutamate prevents the activation loop from adopting the inhibitory conformation. This mutant and activated form of BRAF depends on HSP90 for stability in the cell^15^.

Degradation of proteins by the ubiquitin-proteasome system requires ubiquitylation of lysine residues in the target protein by E3 ubiquitin ligases, which in turn require such substrates to be partially or completely unfolded^18^. Autoinhibited kinases exhibit restrained dynamics^19^, which likely minimizes the exposure of lysine residues for ubiquitylation and consequent degradation by the proteasome. In Src-family kinases, two homologous domains, Src homology 3 and 2 (SH3 and SH2), maintain the kinase domain in an inactive conformation. These interactions “clamp” the kinase and constrain the activation loop in a closed and inhibitory conformation^20,21^. The activation of Src-family kinases proceeds through the dephosphorylation of the phosphotyrosine in the C-terminal tail and the displacement of the SH3-SH2 unit. Whether this clamping mechanism stabilizes kinase domains, making them less reliant on HSP90, is not known.

An interplay of intramolecular interactions and post-translational modifications also underlies autoinhibition in Raf kinases. As shown for BRAF in a quiescent, low-activity state, a single BRAF is doubly phosphorylated on disordered linkers, both N- and C-terminal to the kinase domain. Phosphoresidues in these linkers recruit a 14-3-3 dimer to bind the kinase domain, creating a pocket for the binding of the N-terminal regulatory domains^17,22^. Truncation of the N-terminal segment, including one of the 14-3-3 binding site and cysteine rich domain of BRAF eliminates autoinhibition and the truncated protein exhibits high kinase activity^23,24^.

The HSP90/CDC37 complex can directly bind to and protect unstable client kinases from degradation. Nuclear magnetic resonance studies suggest that the phosphorylated N-terminal segment of CDC37 binds to destabilized kinase domains and recruits HSP90 to these clients^25–27^. If the HSP90 ATPase activity is inhibited pharmacologically, HSP90 cannot load new clients through CDC37 and kinases that are normally HSP90 clients are instead subjected to ubiquitin-mediated degradation^11^. Structures of HSP90 and CDC37 in complex with cyclin-dependent kinase 4^28^, CRAF^29,30^, and BRAF(V600E)^31^ revealed that the chaperone complex binds to a partially unfolded conformation of the kinase domain. Like other chaperones, it is likely that this interaction prevents further unfolding and aggregation of the client kinase domains that would be deleterious to the cell.

We sought to develop a tool to measure the expression levels of kinase molecules in the cell to investigate the effects of mutations on kinase stability. We were inspired by previously reported methods for the use of genetically encoded sensors to study protein turnover through high-throughput screens in live cells^32,33^. We constructed a bicistronic sensor expressing a reference fluorescent protein and a kinase fused to a different fluorescent protein. Normalization of the fluorescence levels of the kinase fusion protein to that of the reference fluorescent protein ensures that measurements are corrected for cell-to-cell variation in levels of transcription or translation. Using this approach, we assessed the relative stability of different protein kinases by measuring their abundance in response to inhibition of HSP90 function or inhibitor binding.

## Results and Discussion

### Design of an Expression-Calibrated Fluorescent Protein Sensor To Monitor Kinase Expression Levels

We developed a lentiviral construct for bicistronic expression of two different, fast maturing, monomeric fluorescent proteins: one fused to a kinase of interest and the other serving as an independent internal control for expression levels. In this system, a spleen focus forming virus (SFFV) promoter^34^ drives expression of mTagBFP2^35^ (a variant of a sea anemone orange fluorescent protein with a maturation time ~14 minutes) followed by the tandem 2A^36,37^ skipping sequence and mNeonGreen^38^ (a variant of an amphioxus yellow/green fluorescent protein with a maturation time ~12 minutes) fused with a long-flexible linker^39^ to a kinase (or any other protein of interest) (figures 1B, detailed schematic in figure S1). For convenience, we refer to mTagBFP2 as BFP and mNeonGreen as GFP in the text and figures.

**Figure 1.**
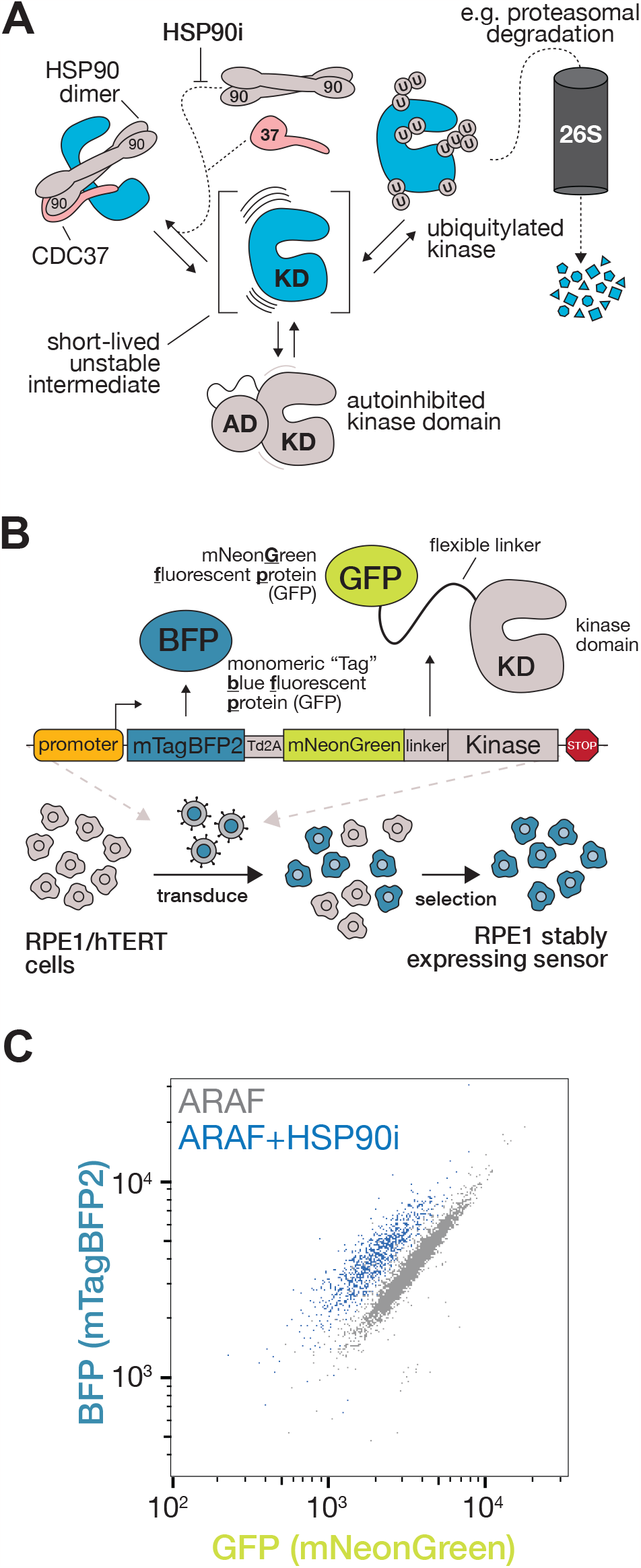
An expression-calibrated, ratiometric fluorescent protein sensor to monitor protein kinase abundance in live cells. A. Schematic of the fates of an unstable kinase domain in the cell. Autoinhibitory domains (AD) often physically stabilize kinase domains by constraining their dynamics. HSP90 and CDC37 collaborate to stabilize a large proportion of kinase domains in the cell, which can be blocked by HSP90 inhbition by small molecules (HSP90i, HSP90 inhibitor). Unstable kinase domains are likely surveilled by ubiquitin ligases in the cell, leading to their proteolytic degradation by the 26S proteasome (26S) when ubiquitylated (U, ubiquitin moieties). B. Schematic of the coding sequence of the lentiviral vector used in this study and the workflow for producing stable cell lines used in this study. A ribosome skipping sequence (tandem 2A, Td2A) allows for BFP (BFP) and GFP (GFP) to be expressed as separate proteins from the same transcript. The GFP sequence is appended to a linker and kinase/kinase domain (KD). The expression cassette is packaged into lentivirus and used to transduce immortalized RPE1 cells expressing human telomerase (hTERT). After transduction, polyclonal populations of cells expressing the sensor can be isolated by FACS. C. Flow cytometry analysis of RPE1/hTERT cells expressing ARAF from the bicistronnic sensor with and without the addition of an HSP90 inhibitor (HSP90i).

Immortalized RPE1 cells expressing human telomerase reverse transcriptase (referred to as RPE1/hTERT or RPE1 cells)^40^ were selected for this study for their robust growth and high lentiviral transduction efficiency. We have also successfully expressed and analyzed the same lentiviral constructs in other cell lines including Ba/F3 murine B lymphocytes, Jurkat T lymphocytes, and HEK 293-derived lines (data not shown). A universal chromatin opening element derived from the human serum response factor promoter^41^ was cloned upstream of the SFFV promoter to mitigate transgene silencing^42^ and ensure robust expression in stable cell lines. The tandem 2A skipping sequence eliminates ribosome readthrough^43^ that could otherwise yield kinases fused to both GFP and BFP. Every molecule of GFP-tagged kinase is linked to the translation of a molecule of BFP, allowing for the ratiometric estimation of GFP-kinase abundance in single cells using fluorescent microscopy or flow cytometry.

Fusion proteins can behave differently than separately expressed proteins that form the fusion protein^44^. To minimize such effects, we attached the kinase to GFP using a long intrinsically disordered linker. We selected an 80 amino acid segment (XTEN80) from a linker designed to adopt a flexible, extended conformation in solution. This linker lacks hydrophobic residues, lysine residues, and has no identifiable short linear motifs^39^. Previous work showed that XTEN80 preserves the enzymatic activity and protein-protein interactions of DNA-modifying enzymes fused to dead Cas9^45^.

### Analysis of HSP90 Protection of Raf Kinases

As a first test of the expression-calibrated sensor, we monitored the levels of Raf-family kinases in cells treated with HSP90 inhibitors. We first cloned full-length, wild-type ARAF fused to GFP downstream of the XTEN80 linker in the construct in figure 1B, generated the corresponding lentivirus, and transduced RPE1/hTERT cells. We then isolated successfully transduced cells by fluorescence activated cell sorting (FACS) of TagBFP2-expressing (BFP+) cells, expanded these cells in culture, and analyzed the expression of BFP and GFP-tagged ARAF by flow cytometry. The selected polyclonal population exhibited a tight correlation between fluorescence intensity of BFP and GFP in each cell (figure 1C), confirming the linked expression of both proteins. When we applied an HSP90 inhibitor (HSP90i) to the cells, the population of cells shifted leftward on the plot, indicating the degradation of GFP tagged ARAF but not BFP (figure 1C).

We cloned full-length, wild-type ARAF, BRAF, and CRAF, as well as full-length BRAF harboring the activating V600E mutation, into the calibrated sensor construct (figure 2A). We applied the HSP90 inhibitor tanespimycin (17-AAG; HSP90i) to transduced cells and analyzed the ratio of GFP fluorescence relative to BFP. As previously shown using different methods, ARAF, CRAF, and BRAF^V600E^ are degraded appreciably after 24 hours when exposed to 1 μM concentration of HSP90i (figure 2D) while wild-type BRAF is unaffected (figure 2D). We determined that this effect was HSP90i dose-dependent and could be recapitulated with other small molecule inhibitors of HSP90, such as geldanamycin and ganetespib (figures 2E, S2D-E).

**Figure 2.**
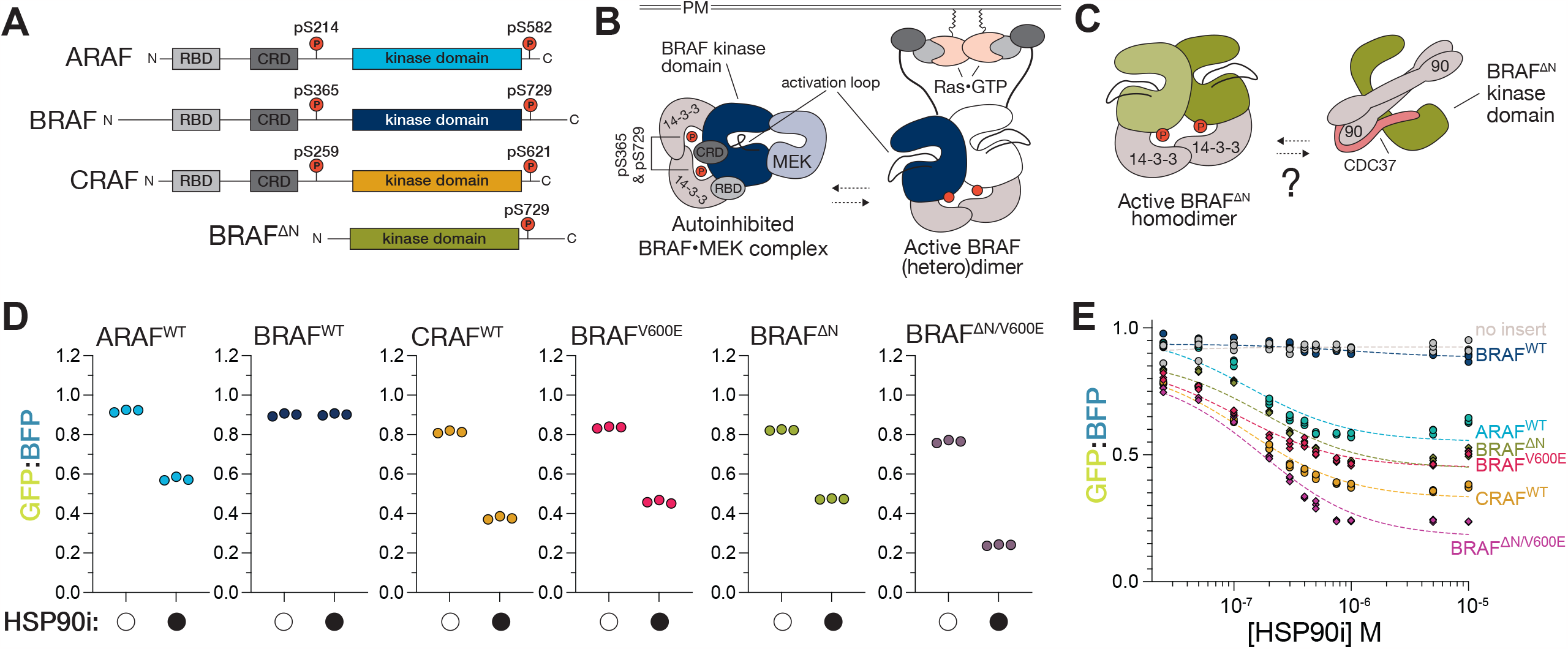
Analysis of HSP90-dependent stability in Raf family kinases. A. Schematic of Raf kinases tested in this experiment. Note the absence of N-terminal regulatory domains in BRAF^ΔN^. (RBD, Ras binding domain; CRD, cysteine rich domain; P, phosphorylation sites). B. A simplified model of BRAF activation in cells. Autoinhibited BRAF bound to MEK and 14-3-3 dimer is recruited to the membrane and activated by GTP-loaded Ras GTPases, dislodging the RBD and CRD from the autoinhibited kinase to promote (homo/hetero)dimerization on a single 14-3-3 dimer. C. Expression of BRAF^ΔN^ in cells yields a highly active dimeric complex in vitro. It is unknown if this active version of BRAF is bound by HSP90/CDC37. D. Plot of levels of Raf kinases and mutants with and without addition of an HSP90i (1 μM tanespimycin), as measured by flow cytometry. (n=3 biological replicates). E. As in 2E, a dose-response plot of normalized levels of Raf kinases and variants. (n=3 biological replicates).

We also analyzed an N-terminal truncation of BRAF (BRAF^ΔN^) lacking its autoinhibitory domains^17^ but retaining other elements required for dimerization^46^ (figure 2C). BRAF^ΔN^ is not autoinhibited and is constitutively dimerized when isolated from cells overexpressing the protein^23,24^. This active form of BRAF is expected to be a client of the HSP90 system. We found that BRAF^ΔN^ and BRAF^V600E^ are degraded similarly in response to HSP90i (figure 2E and F), suggesting active forms of the wild-type kinase BRAF kinase domain are maintained by the HSP90 system. We also introduced the V600E mutation into BRAF^ΔN^ (BRAF^ΔN/V600E^) and observed that this construct is more readily degraded than either BRAF^ΔN^ or BRAF^V600E^ (figures 2E and F, also S2F and G). This result suggests that the N-terminal segment of BRAF may exert some stabilization on the protein even in the context of the V600E-mutant kinase domain.

We evaluated the effects of two other small molecule HSP90 inhibitors, geldanamycin and ganetespib, on Raf kinase expression (figure S2A, D, E). HSP90-client kinases were degraded in response to all three molecules, but ganetespib exhibited superior solubility and stability in culture conditions, and in many cases exhibited a lower IC_50_ in our experiments compared to geldanamycin and tanespimycin. One caveat to the use of ganetespib in our system is that concentrations above 100 nM leads to partial degradation of BRAF^WT^ (figure S1D), which is not the case for the other two inhibitors. This may be due to the slightly different binding preference of ganetespib versus geldanamycin and its derivatives (including 17-AAG) for different conformations of HSP90 that may have variable affinity for different clients^47,48^. Collectively, these studies suggest that there is a fraction of BRAF^WT^ that is stabilized by HSP90 in cells.

### The SH3 and SH2 Domains Contribute to Stabilization of the c-Src Kinase Domain

Src-family kinases consist of a unique N-terminal segment with membrane-targeting motif followed by an SH3, SH2, and kinase domain. The SH3-SH2-kinase unit comprises the well-conserved “Src-module” that is also found in members of the Abl- and Tec-family kinases^49^. In addition to the Src module, the C-terminal segment in Src-family kinases bears a regulatory tyrosine residue important for autoinhibition. A similar SH3-SH2-kinase domain architecture can be found in the Abl- and Tec-family kinases, but they lack the regulatory tyrosine residue in the C-terminal tail. The Src module is an evolutionarily conserved unit that has been repurposed for the regulation of multiple signaling proteins.

When we expressed a c-Src variant corresponding to the Src module followed by the native C-terminal tail (referred to as “c-Src”, figure 4B) embedded in the stability sensor, we found that this species could be expressed to significantly higher levels (~40% increase) compared to the c-Src kinase domain alone (figure 3B). Treatment of both lines with ganetespib resulted in statistically significant but modest reductions in the levels of both proteins (figure 3C). This is consistent with the behavior of endogenous c-Src as a weak HSP90 client reported in previous studies^14,50,51^. These data indicate that the conformational dynamics of the kinase domain is suppressed by the SH2 and SH3 domains^19,52^.

**Figure 3.**
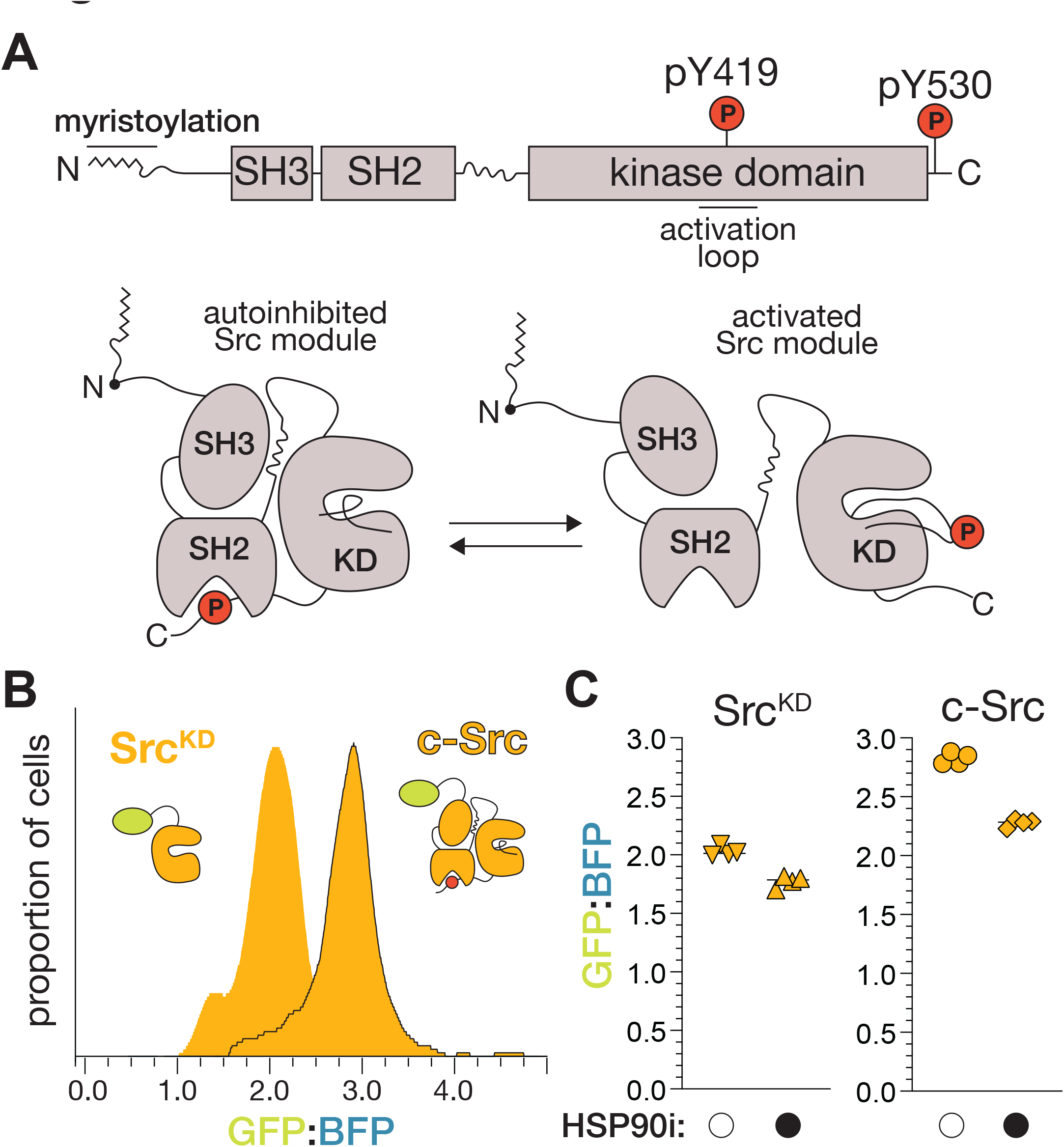
The Src module stabilizes expression of the Src kinase domain. A. Schematic illustrating autoregulation by the Src module. Docking of the SH2 and SH3 domains to the kinase autoinhibit kinase activity in Src-family kinases. B. Expression of Src, is higher when expressed with corresponding SH3, SH2, and tail domains. C. Plots showing both Src^KD^ and C-Src are modestly degraded when cells are treated with HSP90i (ganetespib). Levels of Src^KD^ significantly decreased compared to C-Src in untreaded cells (P<0.01). Groups were analyzed by one-way Brown-Forsythe ANOVA with Dunnett’s T3 multiple comparisons test to generate P-values in the source data file.

**Figure 4.**
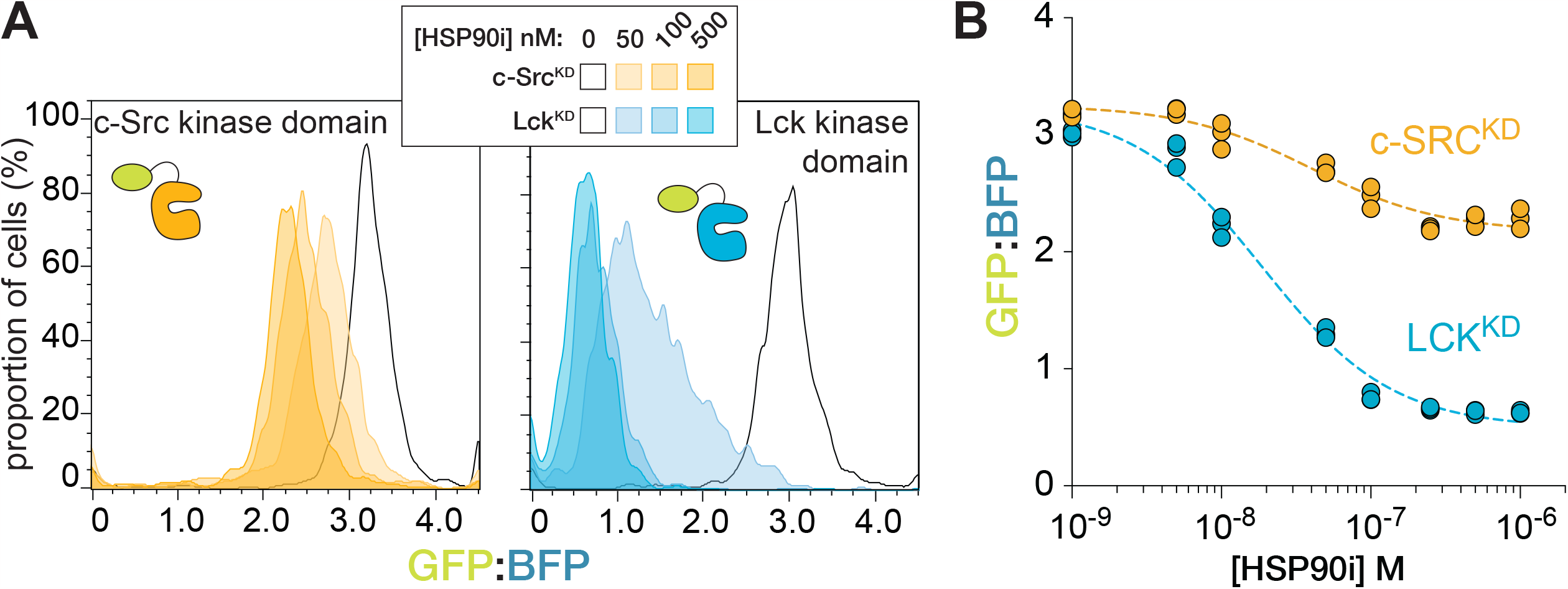
Analysis of HSP90-dependent stability in Src-family kinases. A. Flow cytometry analysis of Src^KD^ and Lck^KD^ in the calibrated sensor in response to varying concentrations of HSP90i (tanespimycin). The outlined curve indicates data from untreated cells. B. Plot of data collected from a similar experiment as in panel 3C demonstrating that Lck^KD^ is degraded to a much greater extent than Src^KD^ in response to HSP90i (tranespimycin).

### Analysis of HSP90-Dependent Stability Across the Src-Family Kinases

We adapted the sensor to study Src-family kinases with their natively modified unique N-terminal segments, to preserve cellular localization and important protein-protein interactions that might affect stability. To design a sensor to accomplish this, we fused kinases at their C-termini to mKOkappa (an orange fluorescent protein, OFP) followed by an independently translated mNeptune2.5 (a far-red fluorescent protein, FRFP) (figure S3B). These fluorescent proteins are spectrally compatible for detection alongside both GFP and BFP, which might facilitate duplexing sensors in the same cell. We compared the behavior of the kinase domain of the strong HSP90 client Lck (Lck^KD^) using both sensors, and found that both are similarly capable of capturing degradation of the kinase after treatment with HSP90i (figure S3A and B).

We introduced seven Src-family kinase domains (c-Src^KD^, Lck^KD^, Fyn^KD^, Lyn^KD^, Blk^KD^, Fgr^KD^, and Yes^KD^) into the calibrated sensor to analyze their HSP90-dependent stability (figure S3C). We found that all seven kinases tested are readily degraded upon HSP90 inhibition, and that this effect does not correlate with the number of lysine residues in each kinase domain (figure S3C and D). Despite this general trend, certain Src-family members, such as Lck, are more reliant on HSP90 to avoid degradation than others^53–56^.

### The Calibrated Sensor Reports on Proteasomal Degradation of HSP90-Client Kinases

We selected c-Src^KD^ and Lck^KD^ as examples of weak and strong HSP90 clients for further characterization. Under basal conditions, the expression of both kinase domains was comparable (figure 4A). However, the Lck kinase domain underwent profound degradation in cells treated with an HSP90i. The majority of Lck^KD^ (>70%) was degraded at submicromolar concentrations of the HSP90 inhibitor while only a small fraction (~20%) of Src^KD^ was destroyed under identical conditions (figure 4B).

To further characterize the sensor constructs, we studied four variants of the sensor: one where GFP is expressed followed solely by the linker sequence (no insert); a second variant in which the well-characterized ornithine decarboxylase degron is appended downstream of the linker (ODC^d2^)^57,58^; and additionally, the previously studied constructs with the c-Src and Lck kinase domains (Src^KD^, Lck^KD^). ODC^d2^ is post-translationally degraded by the ubiquitin proteasome system and should behave similarly to a strong HSP90-client when the chaperone is inhibited. The ODC^d2^ tagged sensor exhibited considerably decreased GFP fluorescence compared to the control lacking any inserted sequence (figure S2A). Inhibition of the proteasome with the small molecule bortezomib partially restored GFP levels in the ODC^d2^ cells (figure S4A). Similarly, the mild proteolytic degradation of Src^KD^ or strong degradation of Lck^KD^ in response to tanespimycin could be reversed by bortezomib in a dose dependent manner (figure S4A).

We tested whether increasing levels of the co-chaperone CDC37 can compensate for the increased kinase degradation observed when HSP90 is inhibited. In vitro studies suggest that CDC37 interacts with kinase domains prior to HSP90 binding^25,27^ and increased CDC37 expression has been shown to compensate for loss of HSP90 activity in yeast^59^. The expression levels of CDC37 are much lower than levels of HSP90 isoforms in mammalian cells, suggesting the CDC37 co-chaperone might be a limiting factor in kinase stabilization^60^. We transduced the four stable cell lines isolated from the previous experiments (no insert, ODC^d2^, c-Src^KD^, and Lck^KD^) with either mCherry alone or wild-type human CDC37 fused to mCherry on its C-terminus. We sorted polyclonal mCherry-expressing populations of cells stably expressing these constructs. We did not observe any appreciable decrease of either c-Src^KD^ or Lck^KD^ degradation in response to HSP90 inhibition regardless of CDC37 overexpression (figure S4C).

Although client binding by HSP90 is thought to be post-translational, some studies have reported that kinases undergo co-translational degradation when the chaperone is inhibited^61,62^. We found that blocking translation with cycloheximide prior to HSP90 inhibition did not reduce the degradation of either c-Src^KD^ or Lck^KD^, suggesting that HSP90-stabilization of c-Src^KD^ and Lck^KD^ from degradation occurs primarily after translation (figure S4B). Kinetic experiments utilizing inhibitors of translation, chaperones, and the proteasome could take advantage of our sensor to comprehensively address how the synthesis, maturation, and degradation of kinases is regulated in live cells. Additionally, replacement of GFP with functionalized enzymes capable of covalent modification, such as HaloTag^63^, could facilitate pulse-chase labeling of client kinases.

### Destabilization of the C-Src Kinase Domain by Activating Mutations

Certain point mutations in full-length c-Src greatly increase its catalytic activity and are also associated with oncogenesis^7^. We sought to determine if such mutations in the kinase domain would alter c-Src^KD^ stability, as measured by our sensor. We studied two point mutations, T341I and E381G (human c-Src numbering; figure 5A).

**Figure 5.**
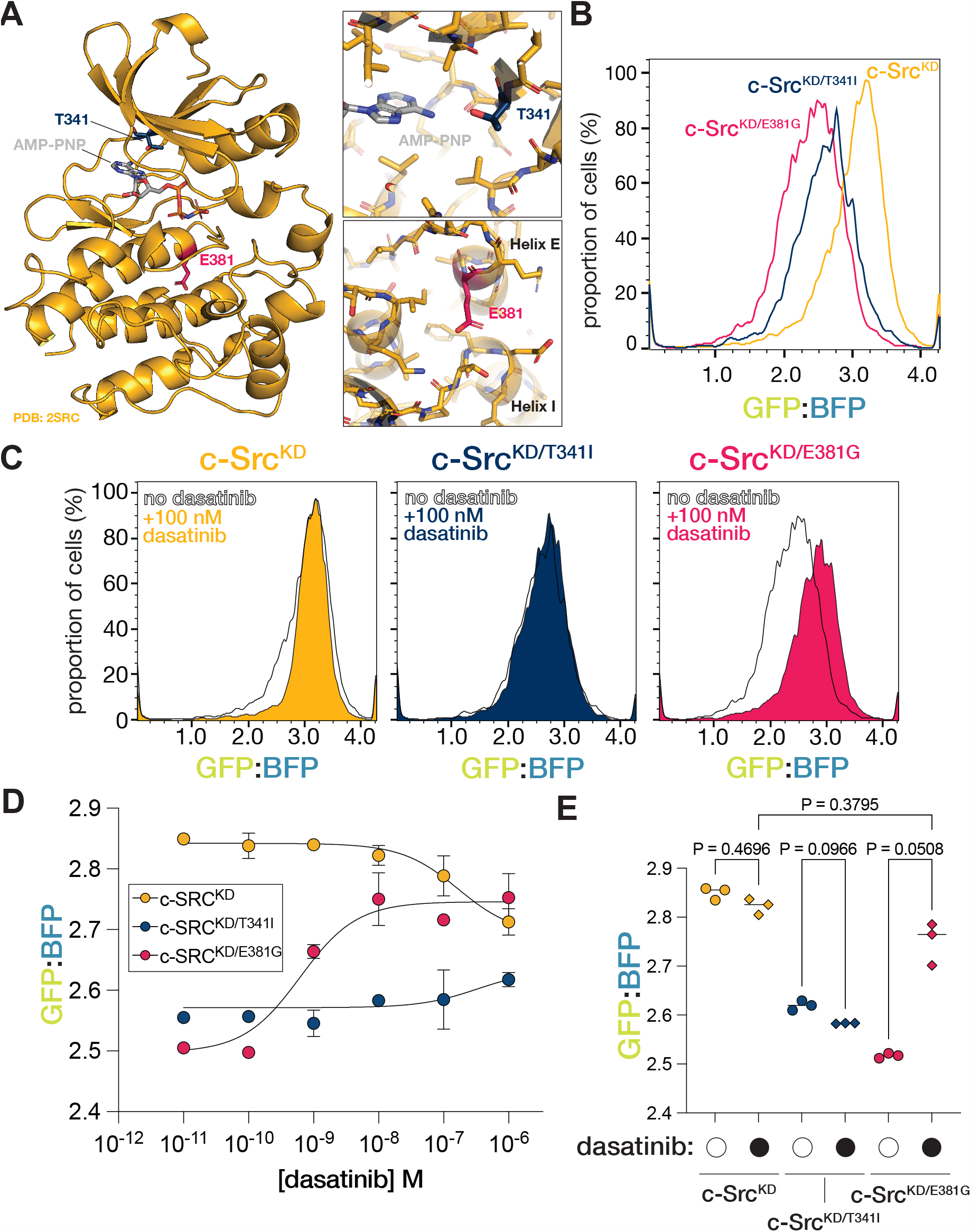
Activating mutations destabilize and a small molecule inhibitor stabilizes the Src kinase domain. A. Illustration of a crystal structure of the Src kinase domain bound to nucleotide. The two residues mutated are shown in the insets. B. Both the T341I and E381G activating mutations destabilize the Src kinase domain as shown by decreased kinase domain levels when expressed from the calibrated sensor. C. Treatment of Src constructs in panel 5B with dasatinib stabilizes the Src^E381G^ but not Src^T341I^ or Src^KD^. D. Dose response curve of Src mutant levels as a function of dasatinib concentration. Error bars represent the standard error of the mean. (n = 3) E. Plot of replicate variant Src^KD^ levels with and without 10 nM dasatinib. Groups were analyzed by one-way Brown-Forsythe ANOVA with Dunnett’s T3 multiple comparisons test to generate P-values on chart.

The T341I mutation was first noted in multiple viral isoforms of chicken c-Src isolated from transformed cells^8,64^. This conserved threonine residue, referred to as the “gatekeeper”, lies within the ATP binding pocket and permits the binding of inhibitors such as imatinib to ABL and gefitinib to EGFR^65,66^. Substitution of the corresponding residue in BCR-ABL or EGFR by bulkier residues, such as isoleucine or methionine, prevents inhibitors from binding and leads to drug resistance^67,68^. Gatekeeper mutations also lead to activation of kinase domains in the absence of inhibitors^65,66^. The E381G mutation was initially identified through screens for mutants of endogenous chicken c-Src that could induce cellular transformation^7^. Both mutations resulted in decreased protein expression levels, suggesting that they destabilize the kinase (figure 5B). The precise mechanism by which these substitutions activate and destabilize c-Src is not well understood^21^.

### Stabilization of Kinase Domains by Kinase Inhibitors

By binding to the kinase domain, kinase inhibitors stabilize certain conformations of their targets^69–71^. Importantly, this action also prevents target kinases from interacting with HSP90/CDC37^72^. Given that the binding of small molecule active-site inhibitors can shield kinase domains from destruction^14,72^, we asked whether the promiscuous tyrosine kinase inhibitor dasatinib can increase levels of mutant c-Src^KD^ (figure 5C). The relative levels of the wild-type c-Src^KD^ did not change in response to the drug. However, treatment of cells expressing c-Src^KD/E381G^ with dasatinib significantly boosted levels of the mutant kinase, comparable to the wild-type kinase in a dose-dependent manner (figure 5D and E). Dasatinib, like other active site inhibitors, does not bind to gatekeeper mutant kinases and has no effect on the levels of c-Src^KD/T341I^.

We extended this approach to study expression levels of BRAF variants. We treated cells expressing BRAF^WT^ (figure S6A) or BRAF^ΔN^ (figure S6B), embedded in our sensor construct, with the inhibitor vemurafenib, which selectively binds the active conformation of the BRAF kinase domain^73^. We observed a subtle but reproducible increase in relative levels of BRAF^ΔN^ in response to high concentrations of vemurafenib (figure S6D). Since the destabilization effects of mutations in BRAF is most apparent when HSP90 is inhibited (figure S2A), we titrated both vemurafenib and the potent HSP90 inhibitor ganetespib. We selected ganetespib as it was the only HSP90 inhibitor tested that could induce mild degradation of BRAF^WT^, thus allowing us to detect any stabilization afforded by vemurafenib. Treatment with high-nanomolar concentrations of the kinase inhibitor vemurafenib protected both BRAF^WT^ and BRAF^ΔN^ from degradation when HSP90 was inhibited by ganetespib (figure S6C and S6D). This suggests that stabilized BRAF kinase domains are resistant to degradation when HSP90 is inhibited.

These experiments demonstrate the capacity of our system to monitor inhibitor-induced stabilization of kinases. Deployment of this tool in high-throughput inhibitor screening could be beneficial for the identification of novel molecules capable of inhibiting mutant kinases as detected by their stabilization under physiologic conditions.

## Conclusions

We developed and validated a fluorescent protein-based tool to measure expression-calibrated protein kinase levels in live mammalian cells. Using this tool, we demonstrated the effects of mutations and small molecules on the stability and degradation of both serine/threonine and tyrosine kinases. We recapitulated previously reported phenomena on the differential reliance of Raf kinases on the HSP90 system, as well as the ability of the V600E substitution to convert BRAF into a strong HSP90 client. Additionally, we explored how autoinhibition of Src-family kinases contributes to their stability, and also showed that the variation within the kinase domains results in different levels of HSP90 dependence. Finally, we demonstrated the ability of the sensor to directly measure the effects of kinase inhibitors on target kinase stability. While calibrated sensors of protein stability have been widely used in the protein quality control field^32,33,74–81^, we show that such tools are also useful in the study of protein kinases.

The stabilization of the kinase domain of c-Src by the SH2 and SH3 domains and C-terminal tail suggests that the kinase domain becomes less stable when released from interactions with the regulatory domains. Given the stabilizing effects of the SH2 and SH3 domains, the stability afforded by the wide range of other autoinhibitory mechanisms in kinases should be explored. For example, the unique myristoylated N-terminus of Abl binds to a hydrophobic pocket in the C-lobe of the kinase domain to facilitate SH2 and SH3 binding to the kinase domain^82–85^. This phenomenon inspired the first clinically successful allosteric kinase inhibitor, asciminib, which blocks BCR-Abl activity by binding to the myristoyl site^86,87^. It is of interest to examine how such an allosteric inhibitor might affect Abl protein stability, as measured by our sensor in cells. Previous studies showed that allosteric inhibitors of BCR-Abl prevent its association with HSP90^14,51^. It is our hope that the calibrated stability sensor that we have developed will be a useful adjunct to future screening campaigns that seek small molecules that bind to and stabilized protein kinases, potentially inhibiting them.

## Methods

### Cell lines and culture

All cell lines were obtained from the UC Berkeley Cell Culture Facility.

HEK 293FT cells were grown in DMEM with high glucose (Gibco, Thermo Fisher #11965092) supplemented with 10% fetal bovine serum (VWR #97068-085), 1x GlutaMAX (Gibco, Thermo Fisher #35050061), 1 mM sodium pyruvate (Gibco, Thermo Fisher #11360070), 1x MEM non-essential amino acids (Gibco, Thermo Fisher #11140050), 10 mM HEPES (Gibco, Thermo Fisher #15630080), and 1x penicillin-streptomycin (Gibco, Thermo Fisher #15140122) and grown at in a humidified 37ºC incubator with 5% atmospheric carbon dioxide.

RPE1/hTERT cells were grown in DMEM/F12 (Gibco, Thermo Fisher #11320033) supplemented with 10% fetal bovine serum (VWR #97068-085), 1x GlutaMAX (Gibco, Thermo Fisher #35050061), 1 mM sodium pyruvate (Gibco, Thermo Fisher #11360070), 1x MEM non-essential amino acids (Gibco, Thermo Fisher #11140050), 10 mM HEPES (Gibco, Thermo Fisher #15630080), and 1x penicillin-streptomycin (Gibco, Thermo Fisher #15140122) on plates pre-coated with 0.1% (w/v) gelatin (Sigma-Aldrich # G9391) in a humidified 37ºC incubator with 5% atmospheric carbon dioxide.

### Lentivirus production and transduction

HEK 293FT cells were plated on six-well or ten-centimeter plates pre-coated with 0.1% (w/v) gelatin (Sigma-Aldrich # G9391). Once cells reached ~90% confluency, they were transfected with 2 μg of lentiviral transfer plasmid, 1.5 μg of psPAX2 (Addgene #12260), and 500 ng of pMD2.G (Addgene #12259) using Lipofectamine 3000 (Invitrogen, Thermo Fisher #L3000001) per the manufacture’s protocol. After 12-18 hours, the media was changed, bovine serum albumin (Cell Signaling Technology #9998) was added to 1% (w/v) along with 2 mM valproic acid (Santa Cruz Biotechnology #sc-202378B) to improve viral gene expression and recovery. After 24 hours, the media was removed from cells and stored at 4ºC overnight and replaced again with DMEM plus BSA and VPA. 24-36 hours later media was again harvested and combined with the previous media and concentrated ~5x by centrifugation using 100 kDa MWCO Amicon Ultra-4 (Millipore # UFC810024) or Amicon Ultra-15 (Millipore # UFC910024) centrifugal filter units at 4ºC. Concentrated virus was applied through a 0.45 μm surfactant free cellulose acetate syringe filter (Nalgene, Fisher Scientific #09-740-106) and immediately used or stored at -80ºC for up to one month. Target cells were transduced in suspension with the filtered virus-containing media in the presence of 8 μg/mL polybrene (Santa Cruz Biotechnology, #sc-134220).

### Drugs and drug treatment

All drugs were purchased from Selleck Chemicals as 10 mM stock solutions in DMSO. Solutions were diluted to twice the final concentration in complete media and negative controls were also made with identical concentrations of DMSO. Cells were plated in 96 well plates in a final volume of 100 μL and incubated overnight for cells to attach to the dish. 100 μL of the corresponding drug dilutions were added to cells the following day and cells were assayed after 36 hours.

### Flow cytometry and fluorescence activated cell sorting

Cells were released from plates with TrypLE (Gibco #12604013) at 37ºC for five to ten minutes. Cells were then diluted in three volumes of flow cytometry buffer (1x PBS, 25 mM HEPES pH 7.4, 1% bovine serum albumin [Cell Signaling Technology #9998], 0.5 mM EDTA, 1x penicillin/streptomycin [Gibco, Thermo Fisher #15140122]) and applied to a 70 μm mesh filter to generate a single cell suspension. Cells were then analyzed on a Thermo Fisher Attune Flow Cytometer and/or sorted on a Sony SH800S cell sorter.

### Data analysis and quantification

The data analyzed to produce dose-response curves and other experiments was derived from the median-fluorescent intensity data collected from of stable lines of cells expressing the calibrated sensor. Biological replicates represent independent transductions and isolates of polyclonal populations of cells expressing the sensor. Statistical analysis of data was conducted in GraphPad Prism.

## Data and Resource Availability

Source data spreadsheet containing all analyzed raw data is attached (sourceData.xlsx). Lentiviral constructs encoding the calibrated sensors will be deposited to Addgene for public distribution.

## Acknowledgements

We would like to thank all members of the Kuriyan lab, Timothy Eisen, Nicholas Ingolia, Shawn Costello, and Natalie Dall for insightful discussions and feedback. We are grateful to Clarise Rivera, Odilia Sianto, Jessica Stewart, and Emily Sumpena for help with experiments. We also thank Mary West of the Cell and Tissue Analysis Facility at UC Berkeley for assistance with equipment and Alison Killilea at the UC Berkeley Cell Culture Facility for reagents and cell lines. This work was supported by a US National Institute of Allergy and Infectious Diseases grant (5P01AI091580-10, J.K.), the Howard Hughes Medical Institute (J.K.), the California Institute of Quantitative Bioscience (J.K.), and Vanderbilt University School of Medicine (J.K.). J.P. was supported by a National Science Foundation Graduate Research Fellowship Award (No. 1752814).

## Author Contributions

J.P. and J.K. conceived and designed the studies. J.P. conducted the experiments. S.M. advised on experiments, suggested the examination of the Src mutations, and helped edit the manuscript. J.W.P. and J.K. analyzed the data and wrote the manuscript.

## Supplemental Figure Legends

Figure S1. Detailed schematic of the lentiviral construct used for calibrated measurements of kinase levels in live cells.

Figure S2. Additional data validating the HSP90-dependence of Raf kinases expressed from the calibrated sensor.

A. Histograms showing the relative GFP/BFP levels from a geldanamycin titration in cells expressing different Raf kinases.

B. Plot of normalized data from Figure 2D.

C. Plot of normalized data from Figure 2D.

D. Comparison of BRAF^WT^ levels in response to titration of three different HSP90 inhibitors.

E. As in panel S1B, but with BRAF^V600E^ expressing cells.

F. Additional biological replicates of the response of different BRAF variants express from the sensor to a titration of tanespimycin.

G. Mean data plotted from panel S1D. (****= p-value < 0.0001, Mann Whitney U Test)

Figure S3. Development and analysis of an orthogonal stability sensor.

A. Schematic of an orthogonal calibrated sensor in which a kinase can be fused to a fluorescent protein (mKOk) on the C-terminus, followed by a skipping sequence and the control fluorescent protein (mNeptune2.5). Left: Example of original sensor expressing the LckKD showing ganetespib (100 nM) promotes degradation of the kinase.

B. As in panel S4C, but with the orthogonal sensor. Note degradation of LckKD in response to ganetespib is robust but with a small dynamic range as compared with LckKD in the original sensor.

C. Schematic of all Src-family kinase domains studied in figure S3D.

D. Levels of different Src-family kinases in response to titration of ganetespib. (data represent mean of n=3 biological replicates)

Figure S4. Characterization of LckKD and SrcKD degradation monitored by the calibrated sensor.

A. Normalized plot of GFP/BFP levels in cells expressing the sensor with no insert, the ODCd2 degron, SrcKD, or LckKD treated with HSP90 inhibitor and/or proteasome inhibitor (bortezomib, BTZ). (n=6 biological replicates).

B. As in panel S2A, but cells were treated with cycloheximide or HSP90 inhibitor. (n=6 biological replicates).

C. Stable expression of mCherry or CDC37-mCherry in sensor expressing cells with and without HSP90i. (n=3 biological replicates).

Figure S5. Histograms of flow cytometry data measuring the effect of HSP90i treatment (tanespimycin, 1 μM) on levels of Src^KD^ variants expressed from the sensor.

Figure S6. Dose-dependent stabilization of BRAF variants by vemurafenib.

A. Dose response curves of BRAF^WT^ treated with or without 1 μM ganetespib and a titration of vemurafenib.

B. Dose response curves of BRAF^ΔN^ treated with or without 1 μM ganetespib and a titration of vemurafenib.

C. Heat map of a two-dimensional titration of vemurafenib and ganetespib in cells expressing BRAF^WT^ from the calibrated sensor. Ratios of the median fluorescent intensity are displayed in each box.

D. As in panel S6E, but with BRAF^ΔN^.

