## supplementalFigures for "A Fluorescence-Based Sensor for Calibrated Measurement of Protein Kinase Stability in Live Cells"

Figure S5. Histograms of flow cytometry data measuring the effect of HSP90i treatment (tanespimycin, 1 $\mu$M) on levels of $Src^{KD}$ variants expressed from the sensor.

Figure S6. Dose-dependent stabilization of BRAF variants by vemurafenib.

A. Dose response curves of $BRAF^{WT}$ treated with or without 1 $\mu$M ganetespib and a titration of vemurafenib.
B. Dose response curves of $BRAF^{\Delta N}$ treated with or without 1 $\mu$M ganetespib and a titration of vemurafenib.
C. Heat map of a two-dimensional titration of vemurafenib and ganetespib in cells expressing $BRAF^{WT}$ from the calibrated sensor. Ratios of the median fluorescent intensity are displayed in each box.

D. As in panel S6E, but with $BRAF^{\Delta N}$.

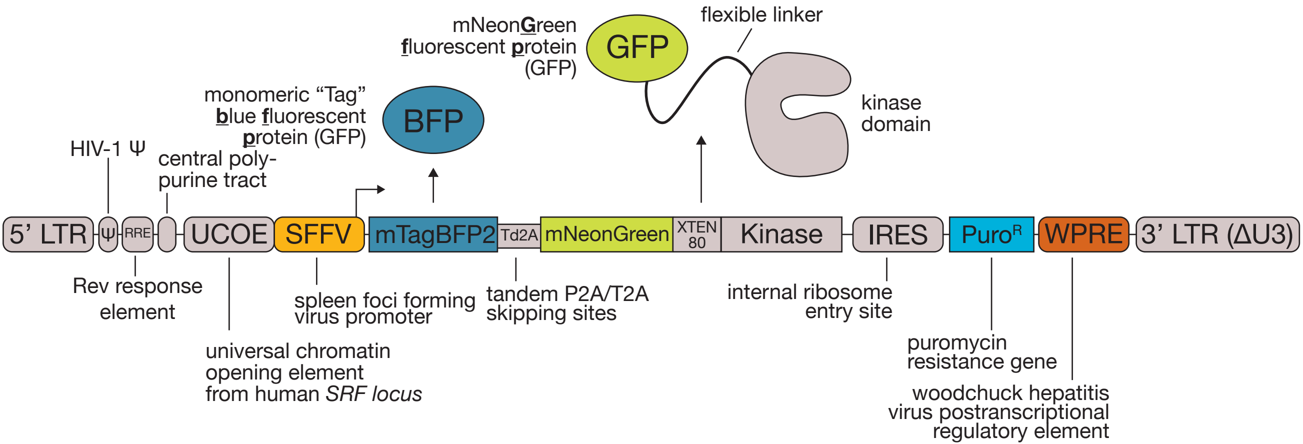

mNeonGreen
fluorescent protein
(GFP)
GFP
flexible linker
kinase
domain
monomeric "Tag"
blue fluorescent
protein (GFP)
BFP
HIV-1 Ψ
central poly-
purine tract
5' LTR
Ψ
RRE
UCOE
SFFV
mTagBFP2
Td2A
mNeonGreen
XTEN
80
Kinase
IRES
Puro^R
WPRE
3' LTR (ΔU3)
Rev response
element
spleen foci forming
virus promoter
tandem P2A/T2A
skipping sites
internal ribosome
entry site
universal chromatin
opening element
from human SRF locus
puromycin
resistance gene
woodchuck hepatitis
virus postranscriptional
regulatory element


### Figure S1

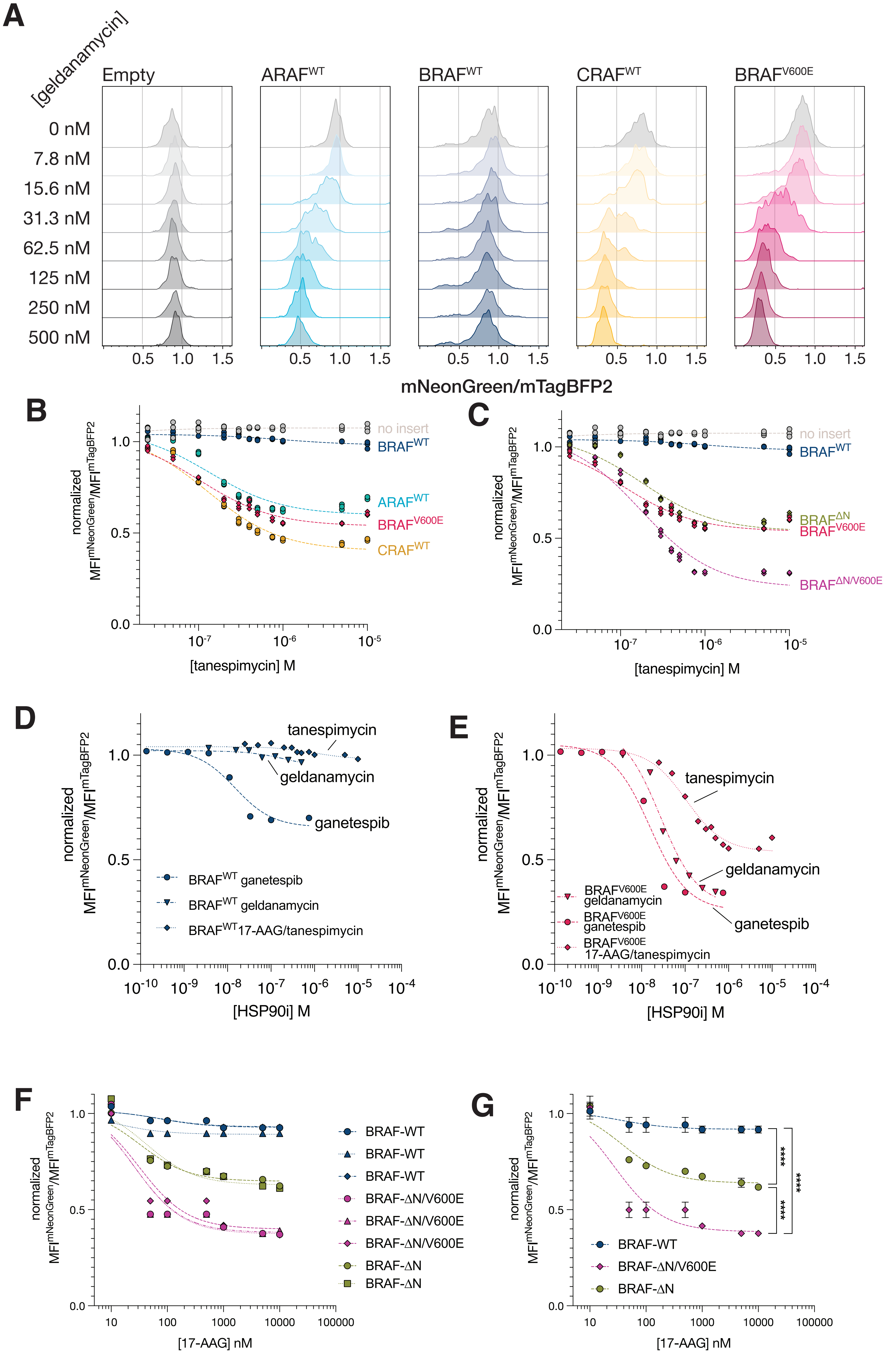

A
[geldanamycin]
Empty
ARAF^WT
BRAF^WT
CRAF^WT
BRAF^V600E
0 nM
7.8 nM
15.6 nM
31.3 nM
62.5 nM
125 nM
250 nM
500 nM
0.5 1.0 1.5
mNeonGreen/mTagBFP2
B
normalized MFI^mNeonGreen/MFI^mTagBFP2
no insert
BRAF^WT
ARAF^WT
BRAF^V600E
CRAF^WT
[tanespimycin] M
C
no insert
BRAF^WT
BRAF^ΔN
BRAF^V600E
BRAF^ΔN/V600E
[tanespimycin] M
D
tanespimycin
geldanamycin
ganetespib
BRAF^WT ganetespib
BRAF^WT geldanamycin
BRAF^WT 17-AAG/tanespimycin
[HSP90i] M
E
tanespimycin
geldanamycin
ganetespib
BRAF^V600E geldanamycin
BRAF^V600E ganetespib
BRAF^V600E 17-AAG/tanespimycin
[HSP90i] M
F
BRAF-WT
BRAF-WT
BRAF-WT
BRAF-ΔN/V600E
BRAF-ΔN/V600E
BRAF-ΔN/V600E
BRAF-ΔN
BRAF-ΔN
[17-AAG] nM
G
****
****
****
BRAF-WT
BRAF-ΔN/V600E
BRAF-ΔN
[17-AAG] nM


Figure S2

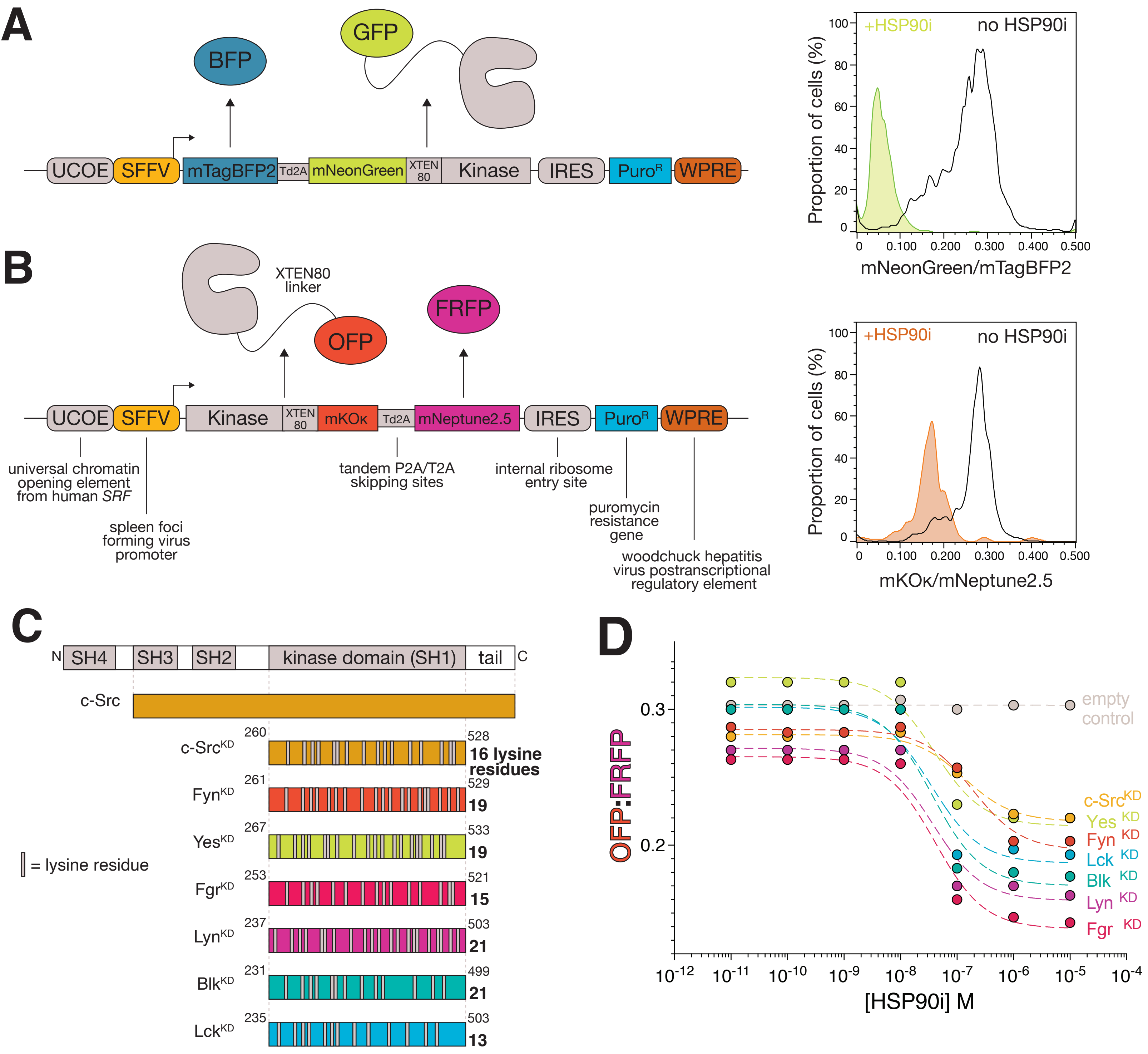
A
BFP
GFP
UCOE
SFFV
mTagBFP2
Td2A
mNeonGreen
XTEN 80
Kinase
IRES
Puro^R
WPRE
+HSP90i
no HSP90i
Proportion of cells (%)
mNeonGreen/mTagBFP2
B
XTEN80 linker
OFP
FRFP
Kinase
XTEN 80
mKOκ
Td2A
mNeptune2.5
universal chromatin opening element from human SRF
spleen foci forming virus promoter
tandem P2A/T2A skipping sites
internal ribosome entry site
puromycin resistance gene
woodchuck hepatitis virus postranscriptional regulatory element
+HSP90i
no HSP90i
Proportion of cells (%)
mKOκ/mNeptune2.5
C
N
SH4
SH3
SH2
kinase domain (SH1)
tail
C
c-Src
c-Src^KD 260 528 16 lysine residues
Fyn^KD 261 529 19
Yes^KD 267 533 19
Fgr^KD 253 521 15
Lyn^KD 237 503 21
Blk^KD 231 499 21
Lck^KD 235 503 13
= lysine residue
D
OFP:FRFP
[HSP90i] M
empty control
c-Src^KD
Yes^KD
Fyn^KD
Lck^KD
Blk^KD
Lyn^KD
Fgr^KD

### Figure S3

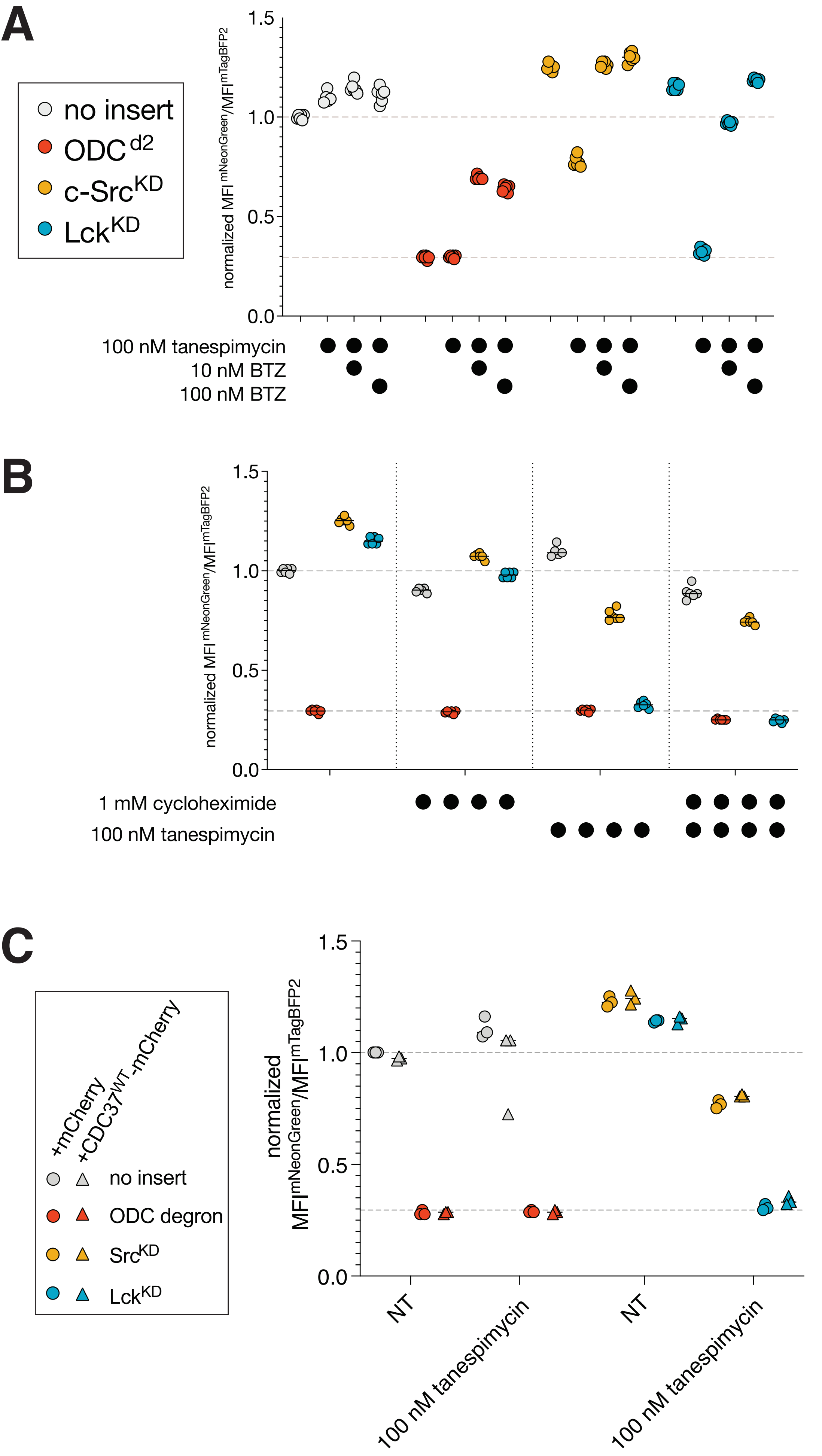
A
no insert
ODC d2
c-Src KD
Lck KD
normalized MFI mNeonGreen/MFI mTagBFP2
1.5
1.0
0.5
0.0
100 nM tanespimycin
10 nM BTZ
100 nM BTZ
B
normalized MFI mNeonGreen/MFI mTagBFP2
1.5
1.0
0.5
0.0
1 mM cycloheximide
100 nM tanespimycin
C
+mCherry
+CDC37 WT -mCherry
no insert
ODC degron
Src KD
Lck KD
normalized MFI mNeonGreen/MFI mTagBFP2
1.5
1.0
0.5
0.0
NT
100 nM tanespimycin
NT
100 nM tanespimycin

Figure S4

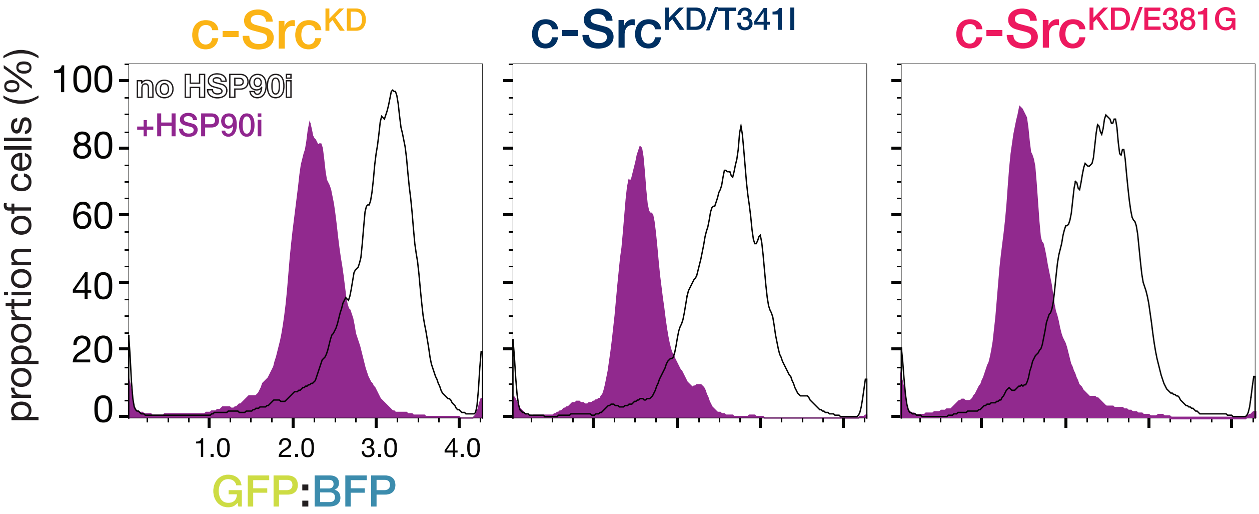

c-SrcKD
c-SrcKD/T341I
c-SrcKD/E381G
no HSP90i
+HSP90i
proportion of cells (%)
100
80
60
40
20
0
1.0
2.0
3.0
4.0
GFP:BFP


### Figure S5

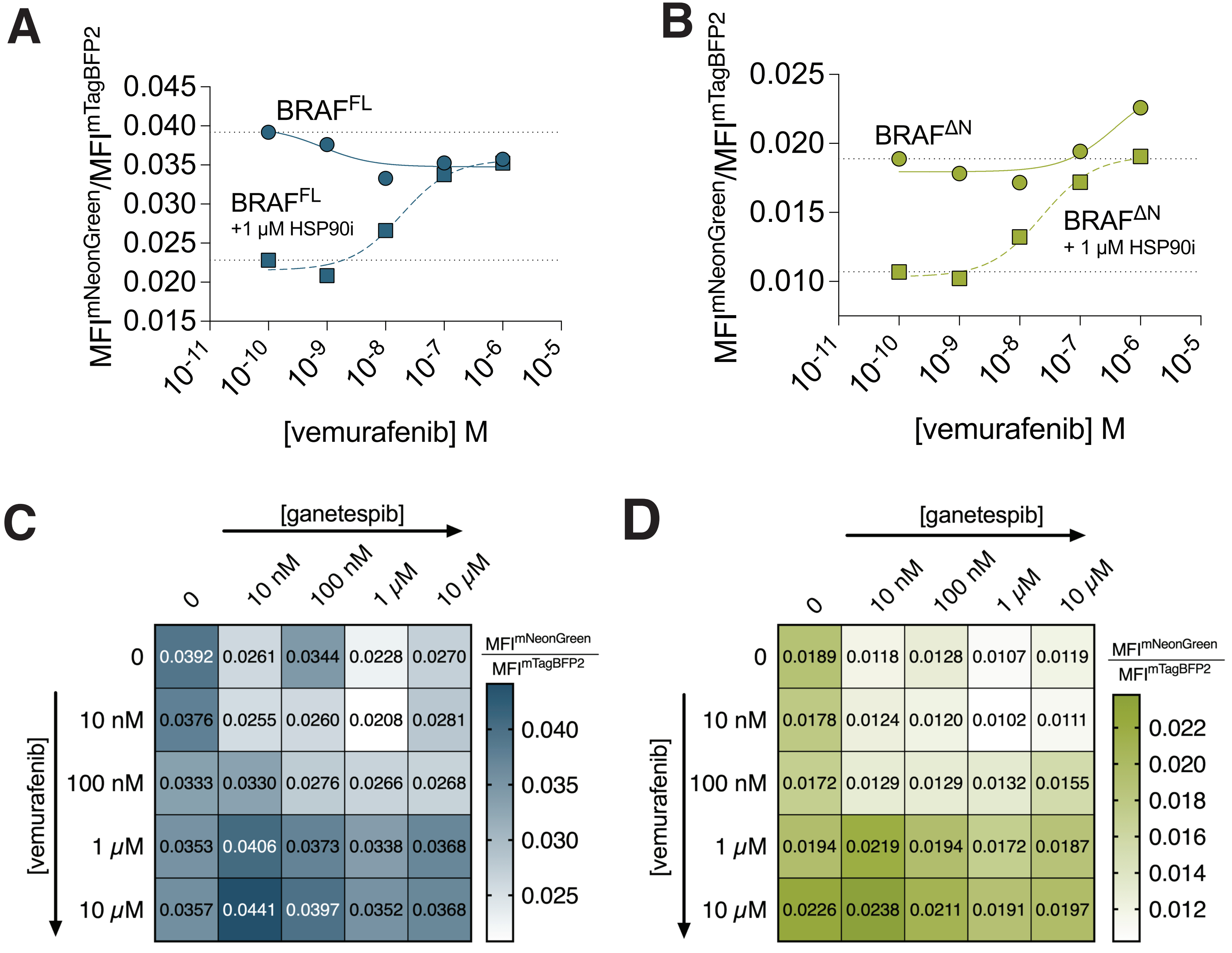
A
MFI^mNeonGreen/MFI^mTagBFP2
0.045
0.040
0.035
0.030
0.025
0.020
0.015
BRAF^FL
BRAF^FL
+1 µM HSP90i
10-11 10-10 10-9 10-8 10-7 10-6 10-5
[vemurafenib] M
B
MFI^mNeonGreen/MFI^mTagBFP2
0.025
0.020
0.015
0.010
BRAF^ΔN
BRAF^ΔN
+ 1 µM HSP90i
10-11 10-10 10-9 10-8 10-7 10-6 10-5
[vemurafenib] M
C
[ganetespib]
0 10 nM 100 nM 1 µM 10 µM
[vemurafenib]
0 0.0392 0.0261 0.0344 0.0228 0.0270
10 nM 0.0376 0.0255 0.0260 0.0208 0.0281
100 nM 0.0333 0.0330 0.0276 0.0266 0.0268
1 µM 0.0353 0.0406 0.0373 0.0338 0.0368
10 µM 0.0357 0.0441 0.0397 0.0352 0.0368
MFI^mNeonGreen / MFI^mTagBFP2
0.040 0.035 0.030 0.025
D
[ganetespib]
0 10 nM 100 nM 1 µM 10 µM
[vemurafenib]
0 0.0189 0.0118 0.0128 0.0107 0.0119
10 nM 0.0178 0.0124 0.0120 0.0102 0.0111
100 nM 0.0172 0.0129 0.0129 0.0132 0.0155
1 µM 0.0194 0.0219 0.0194 0.0172 0.0187
10 µM 0.0226 0.0238 0.0211 0.0191 0.0197
MFI^mNeonGreen / MFI^mTagBFP2
0.022 0.020 0.018 0.016 0.014 0.012

**Figure S6**
